# Selective logging shows no impact on the dietary breadth of the fawn leaf-nosed bat (*Hipposideros cervinus*)

**DOI:** 10.1101/2021.07.30.453964

**Authors:** David R. Hemprich-Bennett, Victoria A. Kemp, Joshua Blackman, Owen T. Lewis, Matthew J. Struebig, Henry Bernard, Stephen J. Rossiter, Elizabeth L. Clare

**Affiliations:** School of Biological and Chemical Sciences, Queen Mary University of London, Mile End Road, London, UK E1 4NS; Department of Zoology, University of Oxford, 11a Mansfield Road, Oxford, UK, OX1 3SZ; Durrell Institute of Conservation and Ecology, University of Kent, Canterbury, Kent, UK, CT2 7NZ; Institute for Tropical Biology and Conservation, Universiti Malaysia Sabah, Kota Kinabalu, Sabah, Malaysia; Department of Biology, York University, 4700 Keele Street, Toronto ON M3J 1P3

## Abstract

Logging activities degrade forest habitats across large areas of the tropics, but the impacts on trophic interactions that underpin forest ecosystems are poorly understood. DNA metabarcoding provides an invaluable tool to investigate such interactions, allowing analysis at a far greater scale and resolution than has previously been possible. We analysed the diet of the insectivorous fawn leaf-nosed bat *Hipposideros cervinus* across a forest disturbance gradient in Borneo, using a dataset of ecological interactions from an unprecedented number of bat-derived faecal samples. Bats predominantly consumed insects from the orders Lepidoptera, Blattodea, Diptera and Coleoptera, and the taxonomic composition of their diet remained relatively consistent across sites regardless of logging disturbance. There was little difference in the richness of prey consumed in each logging treatment, indicating potential resilience of this species to habitat degradation. In fact, bats consumed a high richness of prey items, and intensive sampling is needed to reliably compare feeding ecology over multiple sites regardless of the bioinformatic procedures used.

## Introduction

Logging is a common form of anthropogenic disturbance in forests, with over 90% of those in the tropics logged to some degree (Asner et al., 2009). Most logging undertaken in tropical forests is selective, which tends to favour removal of the largest, and highest-quality trees. While this disturbance can have lasting effects on forest structure (Milodowski et al., 2021) selective logging tends to be much less destructive than clear-felling.

Forest modification through logging is especially pronounced on the island of Borneo, which has lost half of its forest area since 1940 (Gaveau et al., 2014) and 62% of the remaining forest is classified as ‘degraded’ or ‘seriously degraded’ (Gaveau et al., 2016). Most studies of the impact this has on biodiversity have focussed on species composition (e.g. Edwards et al., 2011; Slade et al., 2011; Kitching et al., 2013; Struebig et al., 2013; Deere et al., 2018; Hayward et al., 2021). These often subtle changes to ecological communities can result in changes to ecosystem functioning (Ewers et al., 2015) and the structure of trophic networks (Hemprich-Bennett et al., 2020), indicating that selective logging may alter resilience to future perturbations. Understanding the ecological shifts that take place in degraded forest is of great importance for conservation, especially given the vast scale at which forest is managed for timber extraction globally.

Animal diet can differ between individuals of a species depending on numerous intrinsic and environmental factors. In insectivorous bats for example, inter-individual variation in diet appears to correlate with multiple factors, including wing morphology (Oliveira et al., 2020), sex (Burgar et al., 2014), reproductive condition (Czenze et al., 2018), season (Andriollo et al., 2019; Kolkert et al., 2020), geographic location (Czenze et al., 2018; Vallejo et al., 2019), and habitat (Aizpurua et al., 2018; Hemprich-Bennett et al., 2020; Tournayre et al., 2021). Such variation is of interest when because intraspecific differences in the feeding behaviour of consumers can alter the abundance, community composition and ecological functioning of their prey (Des Roches et al., 2018).

Intraspecific variation in diet is also an important consideration for research design. The analysis of diet in a highly generalist species requires many observations to obtain a representative sample. This can be especially true when studying the dietary ecology of insectivorous bats through metabarcoding, as the technique gives an unprecedented level of taxonomic resolution (Clare et al., 2009), highlighting variation which would not have been apparent with morphological study. Inter-individual variation in bat diet is however often obscured by the use of samples collected from underneath roosts, where numerous bats are defecating (hereafter ‘roost-sourced’ samples) (Clare et al., 2014; Andriollo et al., 2019) and samples cannot be linked to an individual. Obtaining faecal samples from individually identifiable animals (hereafter ‘individual-sourced’ samples) is labour-intensive due to the large trapping effort required, and so while many studies have used individual-sourced samples (e.g. Czenze et al., 2018; Oliveira et al., 2020), their sample sizes tend to be small. Mata et al (2018) used a dataset of individual-sourced samples to analyse the importance of technical and biological replication on the dietary completeness of *Tadarida teniotis* and reiterated the common rule of thumb that 20-50 such samples per species is preferable, but stressed that higher sample sizes may be required for bat species with greater dietary richness or intraspecific variation. The issue of sample size is further complicated in networks generated from metabarcoding data because of methodological considerations such as PCR primer bias and stochasticity (Alberdi et al., 2018), and the influence of bioinformatic choices on the final data analysed (Hemprich-Bennett et al., 2021).

Here we use an unprecedented number of individually-sourced insectivorous bat faecal samples to test the hypothesis that selective logging alters the taxonomic composition and species richness of bats’ diet. We also assess how sample size and bioinformatic parameters affect our inferences of insectivorous diet when using data derived from metabarcoding. Our evaluation focuses on the fawn leaf-nosed bat, *Hipposideros cervinus* - a cave-roosting insectivorous bat found throughout much of maritime Southeast Asia to northeastern Australia. Using high-duty cycle (HDC) echolocation, it is thought to use Doppler-shift compensation to detect the wingbeats of fluttering of prey such as moths (Bell and Fenton, 1984) against a cluttered backdrop (Schnitzler and Kalko, 2001; Lazure and Fenton, 2011). Although some bat species are negatively affected by logging, *H. cervinus* remains a dominant species in both old growth and logged forest in Borneo (Struebig et al., 2013; Hemprich-Bennett et al., 2020). It is not known whether bats such as *H. cervinus* respond to forest degradation by modifying their diets, or are able to maintain stable diets through prey selection or behavioural changes in foraging. We address three main predictions:

1. Taxonomic composition of the diet of *H. cervinus* is altered by rainforest degradation.
2. Individual bats are more specialised in logged forest sites than in primary forest.
3. Estimates of sampling completeness are heavily influenced by MOTU clustering threshold, quality-control methods used and the number of samples.

## Methods

We sampled bats using six harp traps per night at four lowland tropical rainforest sites in Sabah, Malaysia, each <500m above sea level and limited seasonality. Two sites comprise mostly old growth rainforest (Danum Valley and Maliau Basin), and two sites have been subject to substantial logging disturbance (the Sabah Biodiversity Experiment and the Stability of Altered Forest Ecosystems Project) (Supplementary Table 1).

- Old growth rainforest:

- The Danum Valley Conservation Area (hereafter ‘Danum’) is a 438 km^2^ region protected area of old growth rainforest in Sabah (Reynolds et al., 2011). Traps were erected in 2016 for ten nights in a 21-night period and 2017 for ten nights in a 12-night period.
- The Maliau Basin Conservation Area (hereafter ‘Maliau’) is a 588 km2 protected forest made up of lowland and hill forest, most of which has neither been logged nor inhabited in historical times. Traps were erected in 2016 and 2017 for ten nights in a 16-night period.
- Logged forest:

- The Stability of Altered Forest Ecosystems Project (hereafter ‘SAFE’) is a large area of degraded forest being converted to oil palm plantation, with fragments of forest retained for scientific study (Ewers et al., 2011). We sampled in the blocks ‘LFE’, ‘B’ and ‘C’, within the Ulu Segama Forest Reserve and Kalabakan area, during 2015, 2016 and 2017. Each block was sampled for a 5-night period, and then resampled at least 5 weeks later.
- The Sabah Biodiversity Experiment (Hector et al., 2011) (hereafter ‘SBE’) is an area of forest which was logged once in the 1950s and once in the 2000s, and during the sampling period was in the early stages of enrichment replanting (Hector et al., 2011). Sampling took place over a total of 10 nights in a 20-night period in 2016.

Fieldwork, laboratory work and bioinformatics took place as previously described (Hemprich-Bennett et al., 2020). Briefly, bats were captured using harp traps erected along linear features such as streams and trails to target bat flyways. Sampling effort is summarised in Table 1. Faecal samples were processed by DNA extraction, PCR amplification of the CO1 gene using the primers described by Zeale et al (2011), and sequenced on an Illumina MiSeq. For complete methods see (Hemprich-Bennett et al., 2020).

**Table 1.** Trapping effort per site, in harp trap nights. One harp trap night is a harp trap erected for a single night. Six harp traps were used per night, so a single night’s trapping was equal to six harp trap nights.

| **Sample Site** | **2015** | **2016** | **2017** |
| --- | --- | --- | --- |
| **SAFE** | 216 | 180 | 180 |
| **Danum** | 0 | 60 | 60 |
| **Maliau** | 0 | 60 | 60 |
| **SBE** | **0** | **60** | **0** |

### Bioinformatics pipeline

Sequences were assembled into contigs using mothur (Schloss et al., 2009), and forward and reverse primers were removed using the galaxy web platform on the public server at https://usegalaxy.org (Afgan et al., 2016) sequence falling outside of a length of 155-159bp (2bp outside of the expected amplicon length) were excluded from analysis.

When processing the sequence data it is common to cluster sequences into MOTUs (Molecular Operational Taxonomic Units) (Floyd et al., 2002), on the basis of a given threshold of similarity, but the appropriate MOTU clustering thresholds required to best-represent the taxonomic diversity within metabarcoding samples are currently poorly understood (Hemprich‐Bennett et al., 2021). At high clustering thresholds routine sequencing errors may be falsely designated as distinct MOTU, artificially inflating the measured diversity and richness within a sample (Clare et al., 2016). Algorithms implemented using software such as LULU (Frøslev et al., 2017) have been proposed as a method of mitigating this, by combining probable duplicate MOTUs based on patterns of sequence similarity and cooccurrence.

To assess the impact of clustering threshold on the datasets analysed (Hemprich‐Bennett et al., 2021) we generated datasets using MOTU clustering thresholds at ranges 91-98% similarity, using the Uclust algorithm (Edgar, 2010) as implemented in the QIIME platform (Caporaso et al., 2010). Representative sequences for each MOTU per clustering level were then compared to one another using BLAST+ (Camacho et al., 2009), with the resulting data being reduced in LULU (Frøslev et al., 2017) for quality control. All resulting bat-MOTU adjacency lists were then transformed into adjacency matrices using a custom perl script. These matrices were then split into multiple binary adjacency matrices by site. Networks were created by pooling samples from multiple years. To test prediction 2, separate analyses took place on networks both generated as composites of multiple years, and as separate networks for each site and year (see Table 1). All bioinformatic and statistical steps are recorded at https://github.com/hemprichbennett/hice.

### Prediction 1: Taxonomic composition of the diet of *H. cervinus* is altered by rainforest degradation

To analyse the prey taxa consumed by each bat, we used BLAST+ (Camacho et al., 2009) to compare all MOTUs to a library of all arthropod CO1 genes identified to species level using the Barcode of Life Database on 28/03/2018 (BOLD) (Ratnasingham and Hebert, 2007) (3,319,062 sequences), and assigned them taxonomy in MEGAN 6 (Huson et al., 2016) using the parameters in Salinas-Ramos *et al.* (2015). We then assigned MOTUs to order and family level where possible, importing the resulting data into R for analysis, and calculating the proportion of *H. cervinus* individuals per site consuming each taxonomic order. To test the hypothesis that habitat type alters the order-level taxonomic composition of the species’ diet, we analysed the resulting values with a Chi-squared test. The hypothesis was further tested using a permutational multivariate analysis of variance test using distance matrices, and a non-metric multidimensional scaling ordination with 200 permutations using Bray-Curtis dissimilarity, both using the vegan package (Oksanen et al., 2017) on datasets of the order-level diets of each individual bat. We also used a similarity percentages analysis to identify the contribution of each taxonomic order to the observed dissimilarity between sites and years, using Bray-Curtis dissimilarity.

We calculated correlations between the presence/absence of prey orders in faecal samples, using the r package ‘corrplot’ (Wei and Simko, 2017), to identify both potential significant correlations of prey consumption (e.g. bats that feed on Coleoptera may be more likely to feed on Blattodea), and any potential taxonomic bias in PCR.

### Prediction 2: Individual bats are more specialised in logged forest sites than in old growth forest

We created binary bipartite networks for each sampling site and year at 95% similarity clustering and quality control using LULU. In the networks each individual bat and MOTU was classed as a distinct node. A criterion of 95% similarity was chosen for this and all following analyses because it provided a balance between over and under-splitting MOTUs (Hemprich‐Bennett et al., 2021). Using the R package ‘bipartite’ (Dormann, 2011) in R 3.4.4 (R Core Team, 2017) these networks were then analysed using the functions ‘specieslevel’, to calculate the degree of each bat (‘degree’ = the number of prey nodes a bat consumes). Differences between the degree of individuals were compared among sites using an ANOVA with Tukey’s HSD test.

### Prediction 3: Estimates of sampling completeness are heavily influenced by MOTU clustering threshold and quality-control used

Using networks generated at each clustering threshold between 91 and 98% similarity, both with and without quality-control using LULU (Frøslev et al., 2017), we estimated total MOTU richness and sampling completeness of the diet of *H. cervinus* at each site and year using iNEXT (Hsieh et al., 2016), an R package for the interpolation and extrapolation of species diversity using Hill numbers (Chao et al., 2014).

To assess how sample size affects assessments of bat diet, we generated multiple datasets of *n* bats from each site, where *n* was a value of 10-100, increasing in increments of 10 (10, 20, 30, etc), with *n bats* taken at random from each site and the number of MOTUs consumed in that sub-dataset calculated. This was repeated 100,000 times per site and value of *n*, with the resulting data plotted in a violinplot.

## Results

For the full sequencing run of multiple bat species (see Hemprich-Bennett et al., 2020) 18,737,930 contiguous reads were output when assembling the paired-end files. After removing adapters and primers this was reduced to 10,064,815 sequences, which was then further reduced to 932,459 haplotypes after collapsing to haplotype, removing singletons and discarding sequences outside of 2bp of the expected read-length. For full counts of MOTUs before and after clustering with LULU, see Supplementary information 2. Of these, 2,957,444 reads and187,800 haplotypes were derived from *H. cervinus* samples and included in this study.

### Prediction 1: Taxonomic composition of the diet of *H. cervinus* is altered by rainforest degradation

The diet of the bat communities was dominated by insects from the orders Blattodea (especially family Ectobiidae), Diptera (especially family Cecidomyiidae) and Lepidoptera (Figure 1). The chi-squared test showed a non-significant effect of network identity on the order-level composition of a bat populations’ diet (χ^2^ = 0.16, df = 48, p > 0.05). The NMDS showed almost total overlap between the sites (Figure 2) with a stress of 0.21, showing poor convergence. The permutational multivariate analysis of variance test gave an R2 of 0.014 for the explanatory power of site on bat diet. A total of 23 arthropod orders were eaten based on the combined diets of all bats, with Blattodea, Coleoptera, Diptera and Lepidoptera collectively making up at least 79% of all MOTUs identified at each site. Positive correlations were observed between the occurrences of several taxa, with only Araneae and Hymenoptera being negatively correlated with the presence of one another (Supplementary information 3). Blattodea was the only taxon consistently observed to contribute significantly to inter-site dissimilarity scores (SAFE-Maliau p<0.01, SAFE-SBE p = 0.014, Maliau-SBE p = 0.014, SBE-Danum p <0.01, see Supplementary information 4). There was almost complete overlap between the different years sampled at each site (Figure 2) and each site in 2016 (Figure 3).

**Figure 1.**
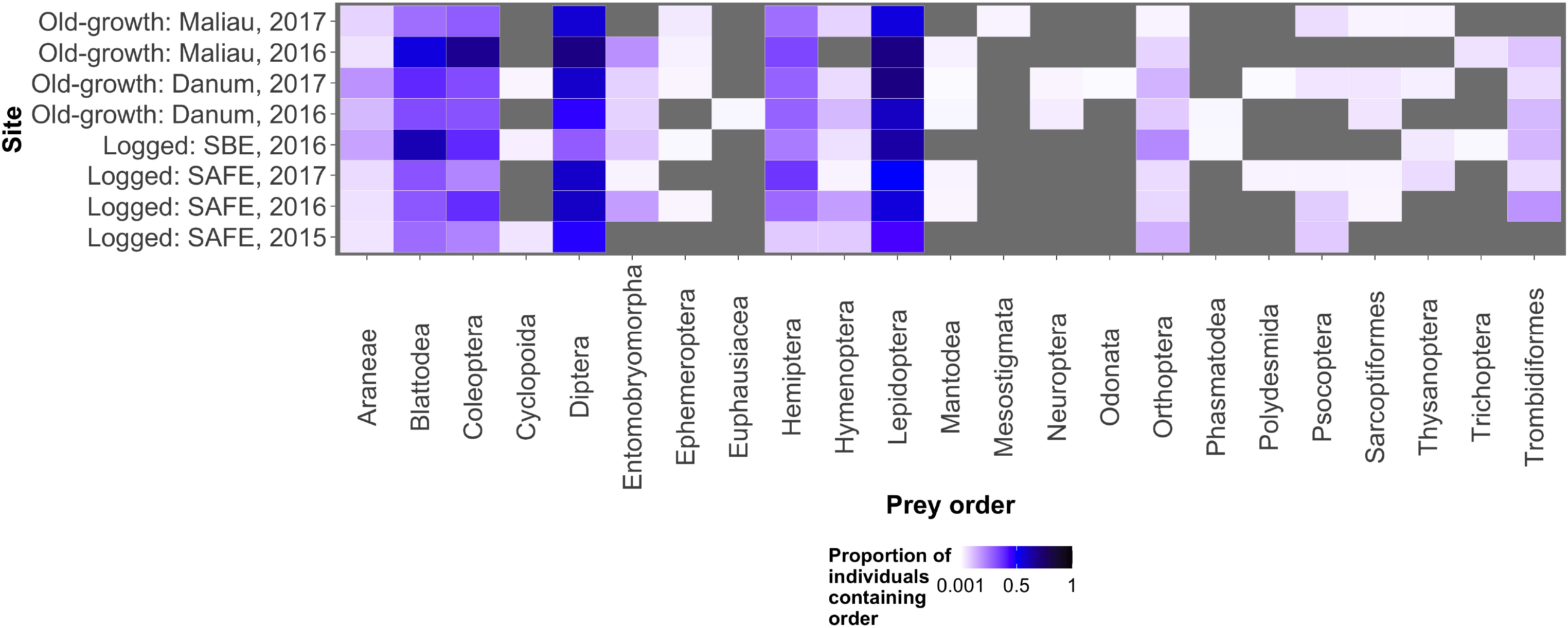
The proportion of all individual bats within a sampling event found to consume each potential prey order. Diptera, Lepidoptera and Blattodea were the commonest prey items, with other prey orders being consumed rarely. The grey background shows locations in the plot where no arthropods of that order were detected in any bats.

**Figure 2.**
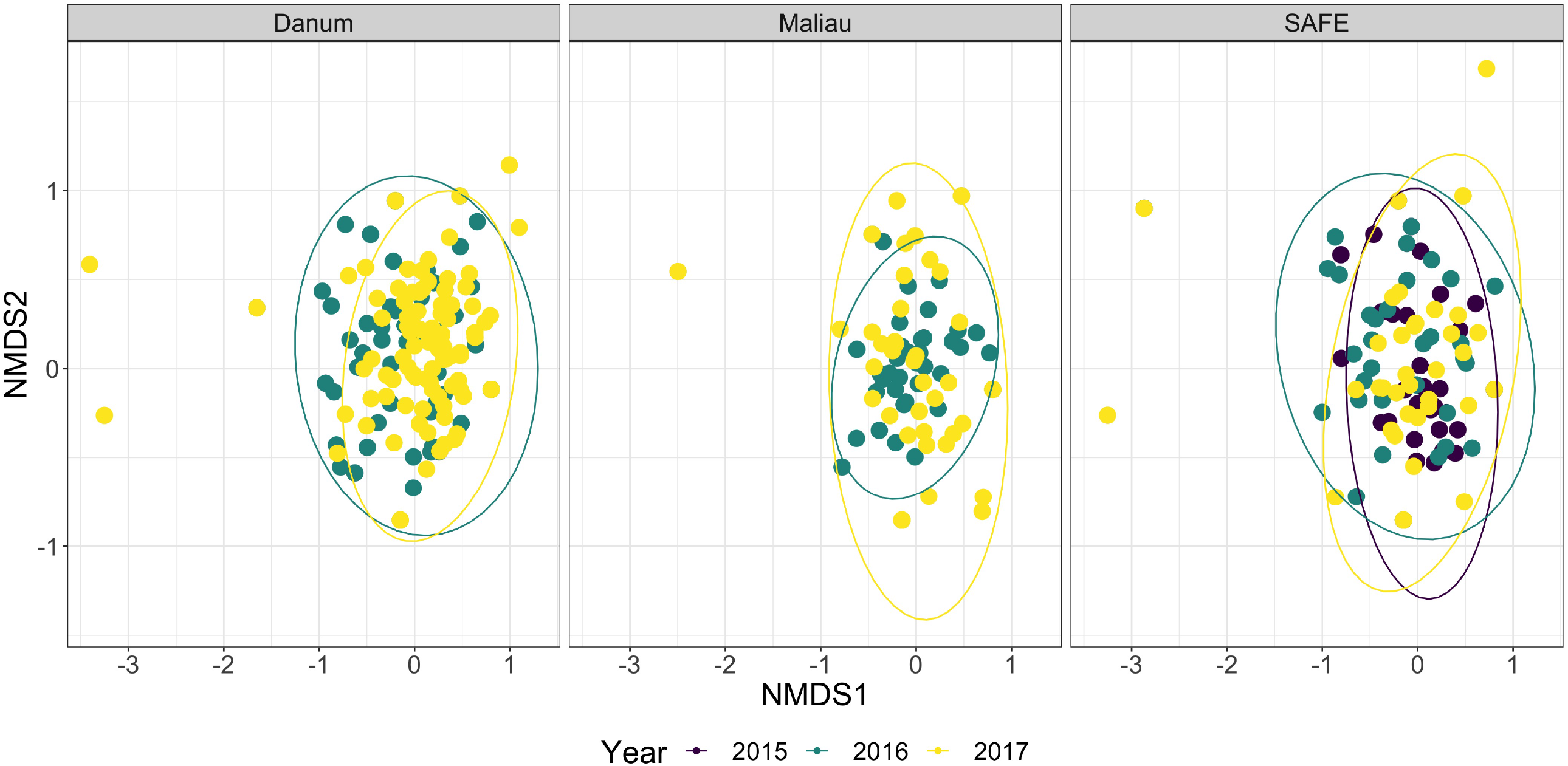
Non-Metric Multidimensional Scaling ordination of the order-level consumption of individual bats across multiple years. The ellipses of each site show almost complete overlap. Stress was 0.21, indicating poor convergence. Danum and Maliau are old-growth sites, SAFE is a logged forest site.

**Figure 3.**
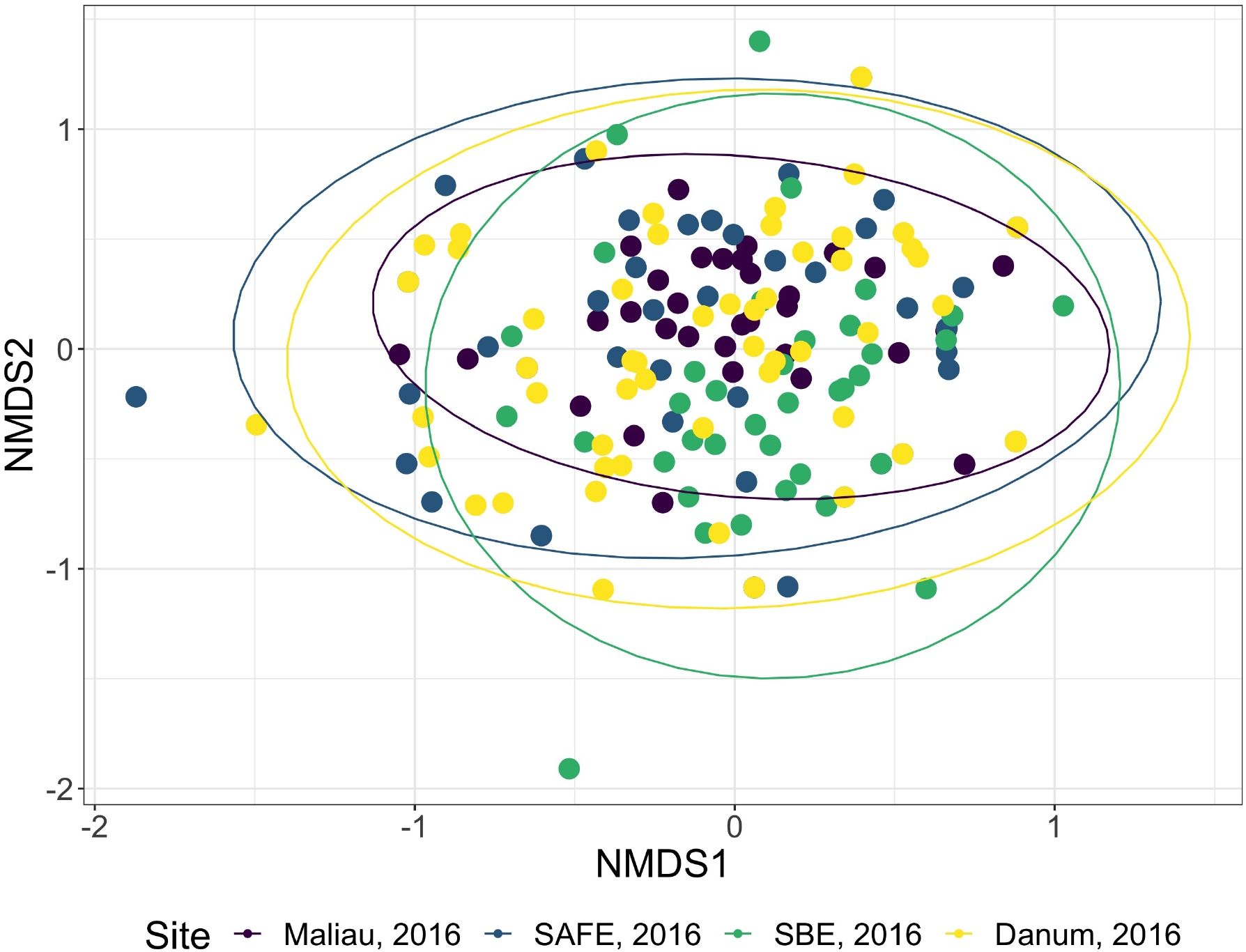
Non-Metric Multidimensional Scaling ordination of the order-level consumption of individual bats in 2016. The ellipses of each site show almost complete overlap. Stress was 0.22, indicating poor convergence. Danum and Maliau are old-growth sites, SAFE and SBE are logged forest sites.

### Prediction 2: Individual bats will be more specialised in logged forest sites than in old growth forest

When comparing networks with all years pooled together, significant differences (p<0.05) were only observed between Danum (old-growth) and SAFE (logged), and between SAFE (logged) and SBE (logged).

### Prediction 3: Estimates of sampling completeness will be heavily influenced by MOTU clustering threshold and quality-control used

None of the networks were estimated as near to fully sampled, with all estimates placing completeness at under 54% (Figure 4), with completeness estimates varying between both sites and years. The number of MOTUs expected increased markedly with clustering threshold when not using LULU for quality control, but this effect was dramatically reduced when using LULU. This algorithm increased estimated sampling completeness by reducing observed and estimated MOTU richness, and lowered the estimated number of samples required to sample the community. Full counts can be found in Supplementary information 2.

**Figure 4.**
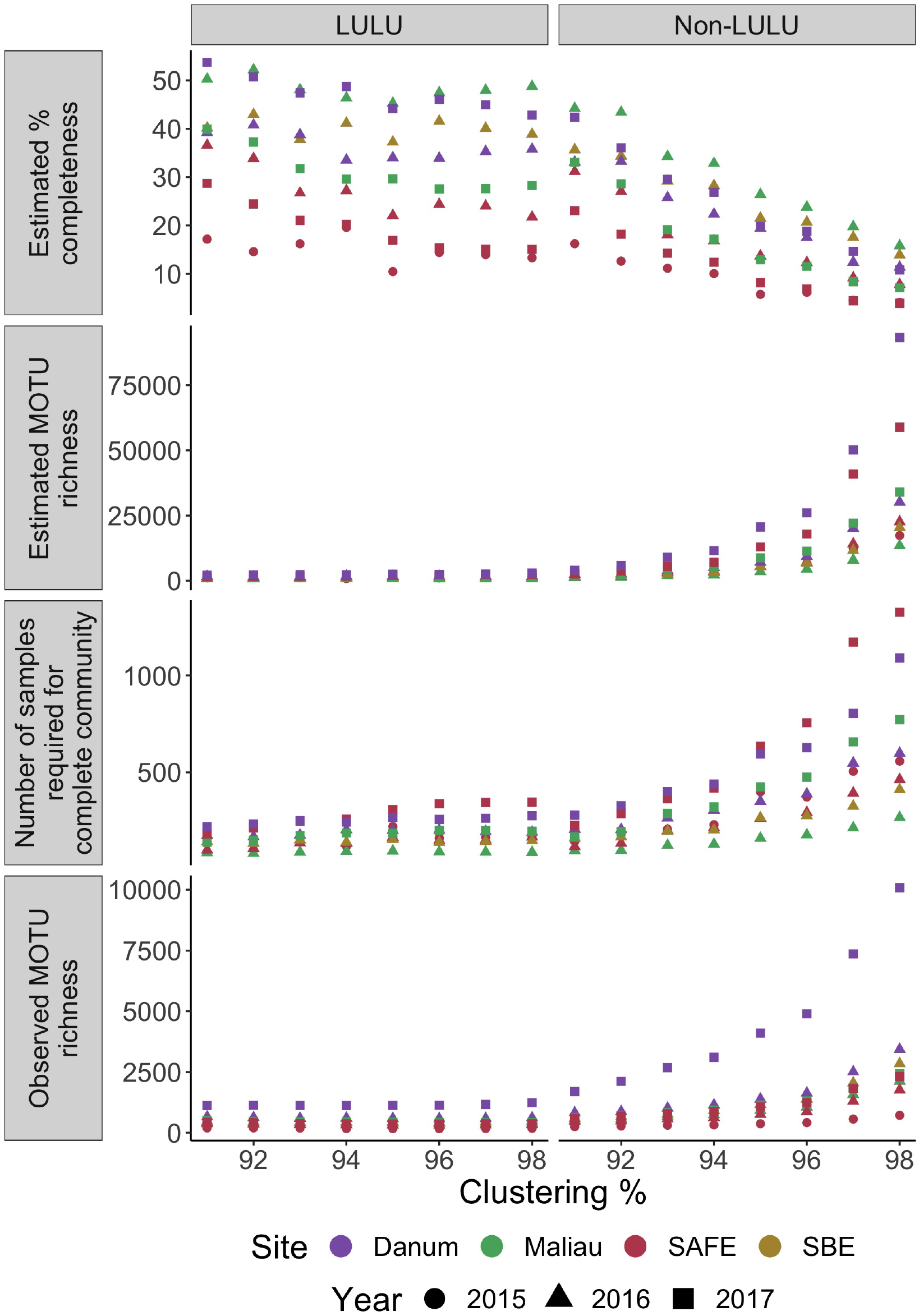
Completeness and richness for each network over a range of MOTU clustering thresholds, with and without use of LULU for post-clustering quality-control. Number of MOTUs is strongly positively correlated with clustering level when not using LULU for quality-control, reducing the estimated completeness of each network.

There was a positive correlation between the number of bats included in a dataset and the number of MOTUs detected (figure 5)

**Figure 5:**
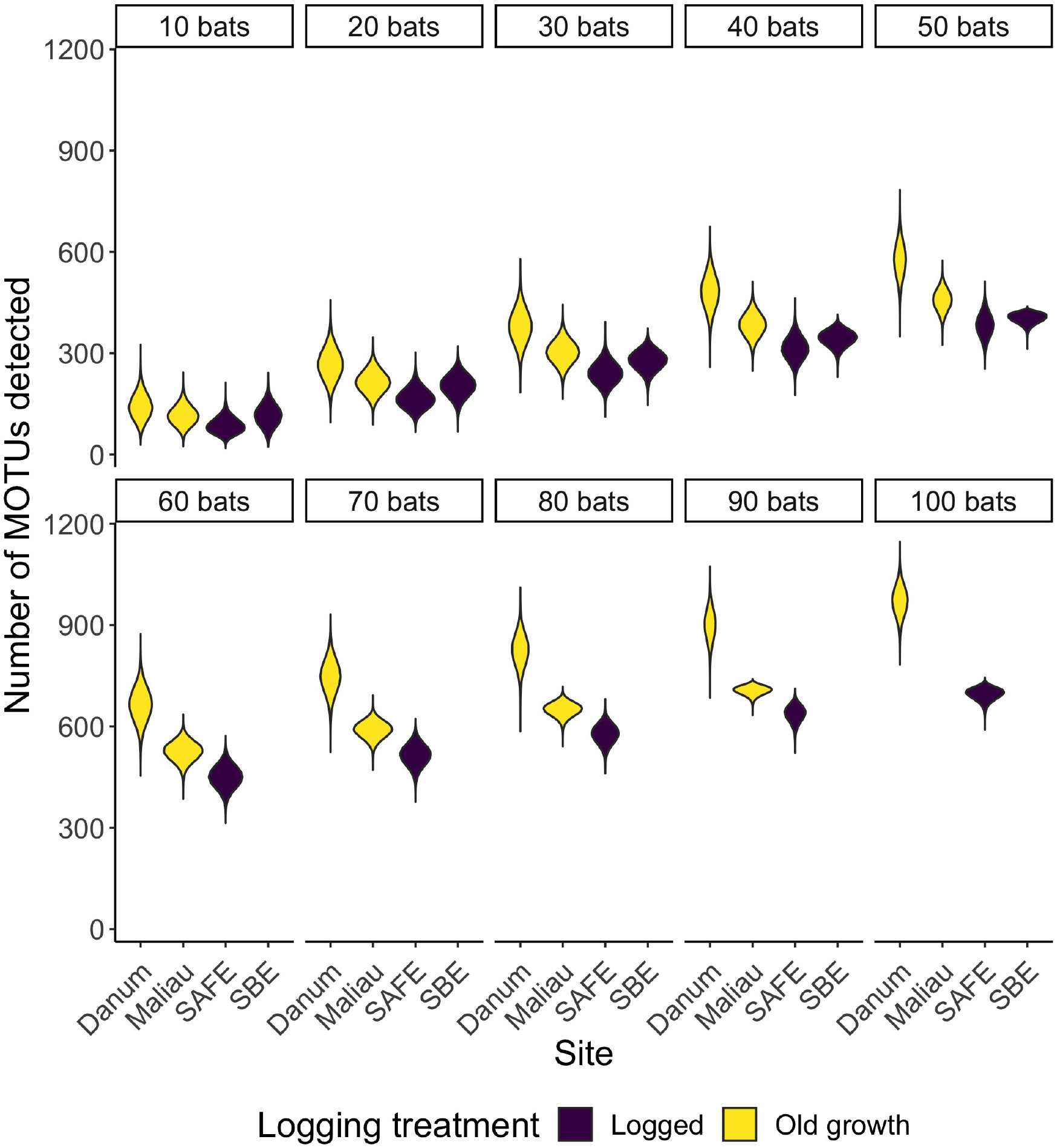
Violinplots showing the distribution of the number of MOTUs consumed when reducing a dataset to *n* bats. With small datasets, sites appear to be rather similar in MOTU richness, but differences emerge as sample sizes increase.

## Discussion

Logging is widespread in tropical forests, yet the consequences of this structural disturbance for trophic interactions are poorly understood. Here we set out to assess how the diet of a generalist insectivorous bat differs between old-growth and degraded forest habitats. We observed broadly similar feeding habits in fawn leaf-nosed bats across forest type with bats consuming many arthropod orders, particularly Blattodea, Coleoptera, Diptera and Lepidoptera. Fawn leaf-nosed bats have extremely high dietary richness, with many hundreds of samples being required to fully capture their diet.

We observed very little alteration in the taxonomic composition of the diet of *H. cervinus*. We saw no significant difference between the consumption of prey at the order-level between sites or years. This suggests that while northeast Borneo may possess high beta-diversity of some insect species (Kitching et al., 2013), at coarse taxonomic levels there is little spatial difference in the prey consumed by *H. cervinus*. Previous findings suggested that, as high-duty cycle echolocators, *H. cervinus* primarily consumed flying insects (Bell and Fenton, 1984; Link et al., 1986; Schnitzler and Kalko, 2001; Lazure and Fenton, 2011), in particular Lepidoptera, Blattodea, Diptera and Coleoptera. The regular presence of diverse families of spiders indicates a dietary contribution of these taxa previously unknown in the Hipposideridae family of bats. Hipposiderids have been observed gleaning stationary targets with fluttering wings (Bell and Fenton, 1984), but the consumption of spiders would either suggest they are gleaning non-fluttering animals, or taking them when ballooning as juveniles. Alternatively, the consumption of spiders could be due to secondary predation: where the bat consumes a primary prey item which has ingested a spider. This seems an unlikely explanation for our dataset, since predatory arthropods other than Araneae are poorly represented in the MOTU dataset. In this study we used one of the most reliable primer sets for amplification of a wide range of digested arthropods (Zeale et al., 2011; Alberdi et al., 2018), but they are also reported to have taxonomic biases towards Diptera and Lepidoptera. However, we found no significant negative correlations between detecting Dipteran or Lepidopteran DNA in a sample, and the detection of any other prey order. This indicates that amplification of dipteran or lepidopteran DNA did not consistently inhibit the amplification of another taxonomic order during PCR, and that sequencing depth is sufficient.

There was no clear pattern of degree differing between logged and old growth habitats. This is in contrast to our previous findings in these study sites (Hemprich-Bennett et al., 2020), that the overall assemblage of bat species in these sites consistently had reduced degree in logged forest than old growth. The diversity of the overall bats’ diet is likely due to the high diversity of prey available to them, and the lack of observed differences in diet between sites may indicate highly flexible foraging, with low impact of land-use change on their diets. Being able to forage adaptively, or fly long distances to viable feeding sites (Struebig et al., 2009) may enable them to remain abundant despite selective logging, while conspecific species experience population declines (Struebig et al., 2013). This species may, as a result, provide ecological redundancy and continue to contribute insectivory when more sensitive bat species have become locally extinct.

A crucial concern in network ecology is the minimum number of samples or observations required to characterise reliably the structure and identity of the interactions within a network (Nielsen and Bascompte, 2007; Rivera-Hutinel et al., 2012). This requirement is complicated in studies utilising DNA metabarcoding as the number of nodes generated is dependent on the bioinformatic choices used to generate them. While MOTU approaches frequently apply a standard resolution to all nodes which helps control for variation in identification, altering MOTU clustering threshold will change the number of nodes and estimates of completeness, analogous to lumping taxonomy-based identifications to higher levels, but without a biological equivalent. We tested MOTU clustering and the use of LULU for quality-control and demonstrated that it was possible to alter estimates of sampling completeness greatly (Figure 4). However, when generating networks with a range of bioinformatics combinations, we observed that none exceeded an estimate of 50% completeness and thus regardless of parameters used, obtaining the full estimate of *H. cervinus* diet would require several hundred samples per site, with the same likely true of many ecologically similar species. Altering MOTU clustering parameters has previously been shown to cause great variation in MOTU counts (Clare et al., 2016) and changes in numerous measures of network-level architecture (Hemprich‐Bennett et al., 2021). The reduction in number of estimated MOTUs provided by LULU (Frøslev et al., 2017) is expected to be of great use in future metabarcoding-based studies to reduce spurious MOTU generation.

The dietary richness found here echoes previous studies (Clare et al., 2009; McCracken et al., 2012) but raises question about the capacity of bats to distinguish between prey types in detail (Neuweiler, 1990) and if this has implications for prey-choice. At the same time, our results highlight the substantial challenge of characterising the diets of this and other insectivorous bat species, especially in hyperdiverse ecosystems such as tropical rainforests. Their large dietary breadth is further highlighted by the fact that DNA extractions performed here were for pooled faecal samples from each individual bat, a technique which Mata *et al.* (Mata et al., 2018) found underestimated the total richness of the diet per bat. Previous intensive studies of arthropod diversity in lowland tropical rainforest have failed to reach an asymptote (Novotný and Basset, 2000; Basset et al., 2012), and if bats are foraging opportunistically it is perhaps unsurprising that the taxonomic breadth of their diet is extremely large and nearly impossible to sample completely.

We demonstrate the vast richness of prey consumed by insectivorous bats in tropical rainforest and show that although quality-control steps in metabarcoding can reduce our estimates of the number of distinct prey items in a site, many hundreds of samples are required to collect a representative description of total diet. Although we focussed our sampling on a single species of insectivorous bat, some inferences likely also apply to similar species, and to other studies that use metabarcoding. The number of sites analysed in this study was low, but it has been shown here that this Hipposiderid species has a highly diverse diet; relying on cockroaches more than previously thought and potentially having a strategy of gleaning non-fluttering prey previously unknown in the family. This bat species is thus thought to exhibit low levels of dietary response to habitat degradation, potentially indicating reasons for their known versatility in the face of landscape modification.

## Acknowledgements

This study was funded by the UK Natural Environment Research Council to SJR, OL and MJS (under the Human-Modified Tropical Forests programme, NE/K016407/1; http://lombok.nerc-hmtf.info/), a Royal Society grant (RG130793) to ELC, and a Bat Conservation International grant to DRHB. We used Queen Mary’s Apocrita HPC facility, supported by QMUL Research-IT (http://doi.org/10.5281/zenodo.438045).

For assistance with data collection we thank Jamiluddin Jami, Arnold James, Mohd. Mustamin, Ampat Siliwong, Sabidee Mohd. Rizan, Najmuddin Jamal, Genevieve Durocher and Anne Seltmann. We thank the Sabah Biodiversity Council, Sabah Forest Department, Yayasan Sabah, and Benta Wawasan Sdn. Bhd. for research permissions (Access licenses: JKM/MBS.1000-2/2 (374), JKM/MBS.1000-2/2 JLD.4 (23), JKM/MBS.1000-2/2 JLD.4 (45), JKM/MBS.1000-2/2 JLD.4 (41), JKM/MBS.1000-2/2 JLD.4 (46), JKM/MBS.1000-2/2 JLD.5 (123), JKM/MBS.1000-2/2 JLD.5 (153), Export licenses: JKM/MBS.1000-2/3 JLD.2(55), JKM/MBS.1000-2/3 JLD.2 (95), JKM/MBS.1000-2/3 JLD.3 (31)) We thank Eleanor Slade and members of the LOMBOK consortium for facilitating research in Sabah, and we are grateful to the Sabah Biodiversity Council (Danum Valley access permits: YS/DVMC/2015/221, YS/DVMC/2016/11, YS/DVMC/2015/222, YS/DVMC/2016/13, YS/DVMC/2017/42, YS/DVMC/2017/41, Maliau Basin access permits: YS/MBMC/2015/186, YS/MBMC/2016/23, YS/MBMC/2015/187, YS/MBMC/2016/25, YS/MBMC/2017/67, YS/MBMC/2017/66)).

We thank Steven Le Comber, Hernani Oliveira, Joshua Potter, Sandra Álvarez Carretero and Kim Warren for their analytical assistance, and Mark Brown and Darren Evans, for helpful comments on earlier versions of this manuscript.

## Works cited

Afgan, E., Baker, D., van den Beek, M., Blankenberg, D., Bouvier, D., Čech, M., et al. (2016). The Galaxy platform for accessible, reproducible and collaborative biomedical analyses: 2016 update. Nucleic Acids Res. 44, W3–W10. doi:10.1093/nar/gkw343.

Aizpurua, O., Budinski, I., Georgiakakis, P., Gopalakrishnan, S., Ibañez, C., Mata, V., et al. (2018). Agriculture shapes the trophic niche of a bat preying on multiple pest arthropods across Europe: Evidence from DNA metabarcoding. Mol. Ecol. 27, 815–825. doi:https://doi.org/10.1111/mec.14474.

Alberdi, A., Aizpurua, O., Gilbert, M. T. P., and Bohmann, K. (2018). Scrutinizing key steps for reliable metabarcoding of environmental samples. Methods Ecol. Evol. 9, 134–147. doi:10.1111/2041-210X.12849.

Andriollo, T., Gillet, F., Michaux, J. R., and Ruedi, M. (2019). The menu varies with metabarcoding practices: A case study with the bat *Plecotus auritus*. PLOS ONE 14, e0219135. doi:10.1371/journal.pone.0219135.

Asner, G. P., Rudel, T. K., Aide, T. M., Defries, R., and Emerson, R. (2009). A Contemporary Assessment of Change in Humid Tropical Forests. Conserv. Biol. 23, 1386–1395. doi:https://doi.org/10.1111/j.1523-1739.2009.01333.x.

Basset, Y., Cizek, L., Cuénoud, P., Didham, R. K., Guilhaumon, F., Missa, O., et al. (2012). Arthropod Diversity in a Tropical Forest. Science 338, 1481–1484. doi:10.1126/science.1226727.

Bell, G. P., and Fenton, M. B. (1984). The use of Doppler-shifted echoes as a flutter detection and clutter rejection system: the echolocation and feeding behavior of *Hipposideros ruber* (Chiroptera: Hipposideridae). Behav. Ecol. Sociobiol. 15, 109–114. doi:10.1007/BF00299377.

Burgar, J. M., Murray, D. C., Craig, M. D., Haile, J., Houston, J., Stokes, V., et al. (2014). Who’s for dinner? High-throughput sequencing reveals bat dietary differentiation in a biodiversity hotspot where prey taxonomy is largely undescribed. Mol. Ecol. 23, 3605–3617. doi:10.1111/mec.12531.

Camacho, C., Coulouris, G., Avagyan, V., Ma, N., Papadopoulos, J., Bealer, K., et al. (2009). BLAST+: architecture and applications. BMC Bioinformatics 10, 421. doi:10.1186/1471-2105-10-421.

Caporaso, J. G., Kuczynski, J., Stombaugh, J., Bittinger, K., and Bushman, F. D. (2010). QIIME allows analysis of high-throughput community sequencing data. Nat. Methods 7, 335–336. doi:10.1038/nmeth.f.303.

Chao, A., Gotelli, N. J., Hsieh, T. C., Sande, E. L., Ma, K. H., Colwell, R. K., et al. (2014). Rarefaction and extrapolation with Hill numbers: a framework for sampling and estimation in species diversity studies. Ecol. Monogr. 84, 45–67.

Clare, E. L., Chain, F. J. J., Littlefair, J. E., and Cristescu, M. E. (2016). The effects of parameter choice on defining molecular operational taxonomic units and resulting ecological analyses of metabarcoding data. Genome 59, 981–990. doi:10.1139/gen-2015-0184.

Clare, E. L., Fraser, E. E., Braid, H. E., Fenton, M. B., and Hebert, P. D. N. (2009). Species on the menu of a generalist predator, the eastern red bat *Lasiurus borealis*: using a molecular approach to detect arthropod prey. Mol. Ecol. 18, 2532–2542. doi:10.1111/j.1365-294X.2009.04184.x.

Clare, E. L., Symondson, W. O. C., Broders, H., Fabianek, F., Fraser, E. E., MacKenzie, A., et al. (2014). The diet of *Myotis lucifugus* across Canada: assessing foraging quality and diet variability. Mol. Ecol. 23, 3618–3632. doi:10.1111/mec.12542.

Czenze, Z. J., Tucker, J. L., Clare, E. L., Littlefair, J. E., Hemprich‐Bennett, D. R., Oliveira, H. F. M., et al. (2018). Spatiotemporal and demographic variation in the diet of New Zealand lesser short-tailed bats (*Mystacina tuberculata*). Ecol. Evol. 8, 7599–7610. doi:10.1002/ece3.4268.

Deere, N. J., Guillera‐Arroita, G., Baking, E. L., Bernard, H., Pfeifer, M., Reynolds, G., et al. (2018). High Carbon Stock forests provide co-benefits for tropical biodiversity. J. Appl. Ecol. 55, 997–1008. doi:https://doi.org/10.1111/1365-2664.13023.

Des Roches, S., Post, D. M., Turley, N. E., Bailey, J. K., Hendry, A. P., Kinnison, M. T., et al. (2018). The ecological importance of intraspecific variation. Nat. Ecol. Evol. 2, 57–64. doi:10.1038/s41559-017-0402-5.

Dormann, C. F. (2011). How to be a specialist? Quantifying specialisation in pollination networks. Netw. Biol. 1, 1–20.

Edgar, R. C. (2010). Search and clustering orders of magnitude faster than BLAST. Bioinformatics 26, 2460–2461. doi:10.1093/bioinformatics/btq461.

Edwards, D. P., Larsen, T. H., Docherty, T. D. S., Ansell, F. A., Hsu, W. W., Derhé, M. A., et al. (2011). Degraded lands worth protecting: the biological importance of Southeast Asia’s repeatedly logged forests. Proc. R. Soc. B Biol. Sci. 278, 82–90. doi:10.1098/rspb.2010.1062.

Ewers, R. M., Boyle, M. J. W., Gleave, R. A., Plowman, N. S., Benedick, S., Bernard, H., et al. (2015). Logging cuts the functional importance of invertebrates in tropical rainforest. Nat. Commun. 6, 6836. doi:10.1038/ncomms7836.

Ewers, R. M., Didham, R. K., Fahrig, L., Ferraz, G., Hector, A., Holt, R. D., et al. (2011). A large-scale forest fragmentation experiment: the Stability of Altered Forest Ecosystems Project. Philos. Trans. R. Soc. B Biol. Sci. 366, 3292–3302. doi:10.1098/rstb.2011.0049.

Floyd, R., Abebe, E., Papert, A., and Blaxter, M. (2002). Molecular barcodes for soil nematode identification. Mol. Ecol. 11, 839–850. doi:10.1046/j.1365-294X.2002.01485.x.

Frøslev, T. G., Kjøller, R., Bruun, H. H., Ejrnæs, R., Brunbjerg, A. K., Pietroni, C., et al. (2017). Algorithm for post-clustering curation of DNA amplicon data yields reliable biodiversity estimates. Nat. Commun. 8, 1188. doi:10.1038/s41467-017-01312-x.

Gaveau, D. L. A., Sheil, D., Husnayaen Salim, M. A., Arjasakusuma, S., Ancrenaz, M., et al. (2016). Rapid conversions and avoided deforestation: examining four decades of industrial plantation expansion in Borneo. Sci. Rep. 6, 32017. doi:10.1038/srep32017.

Gaveau, D. L. A., Sloan, S., Molidena, E., Yaen, H., Sheil, D., Abram, N. K., et al. (2014). Four Decades of Forest Persistence, Clearance and Logging on Borneo. PLOS ONE 9, e101654. doi:10.1371/journal.pone.0101654.

Hayward, R. M., Banin, L. F., Burslem, D. F. R. P., Chapman, D. S., Philipson, C. D., Cutler, M. E. J., et al. (2021). Three decades of post-logging tree community recovery in naturally regenerating and actively restored dipterocarp forest in Borneo. For. Ecol. Manag. 488, 119036. doi:10.1016/j.foreco.2021.119036.

Hector, A., Philipson, C., Saner, P., Chamagne, J., Dzulkifli, D., O’Brien, M., et al. (2011). The Sabah Biodiversity Experiment: a long-term test of the role of tree diversity in restoring tropical forest structure and functioning. Philos. Trans. R. Soc. Lond. B Biol. Sci. 366, 3303–3315. doi:10.1098/rstb.2011.0094.

Hemprich-Bennett, D. R., Kemp, V. A., Blackman, J., Struebig, M. J., Lewis, O. T., Rossiter, S. J., et al. (2020). Altered structure and stability of bat-prey interaction networks in logged tropical forests revealed by metabarcoding. bioRxiv, 2020.03.20.000331. doi:10.1101/2020.03.20.000331.

Hemprich‐Bennett, D. R., Oliveira, H. F. M., Comber, S. C. L., Rossiter, S. J., and Clare, E. L. (2021). Assessing the impact of taxon resolution on network structure. Ecology 102, e03256. doi:https://doi.org/10.1002/ecy.3256.

Hsieh, T. C., Ma, K. H., and Chao, A. (2016). iNEXT: Interpolation and Extrapolation for Species Diversity. Available at: http://chao.stat.nthu.edu.tw/blog/software-download/.

Huson, D. H., Beier, S., Flade, I., Górska, A., El-Hadidi, M., Mitra, S., et al. (2016). MEGAN Community Edition - Interactive Exploration and Analysis of Large-Scale Microbiome Sequencing Data. PLOS Comput. Biol. 12, e1004957. doi:10.1371/journal.pcbi.1004957.

Kitching, R. L., Ashton, L. A., Nakamura, A., Whitaker, T., and Khen, C. V. (2013). Distance-driven species turnover in Bornean rainforests: homogeneity and heterogeneity in primary and post-logging forests. Ecography 36, 675–682. doi:10.1111/j.1600-0587.2012.00023.x.

Kolkert, H., Andrew, R., Smith, R., Rader, R., and Reid, N. (2020). Insectivorous bats selectively source moths and eat mostly pest insects on dryland and irrigated cotton farms. Ecol. Evol. 10, 371–388. doi:https://doi.org/10.1002/ece3.5901.

Lazure, L., and Fenton, M. B. (2011). High duty cycle echolocation and prey detection by bats. J. Exp. Biol. 214, 1131–1137. doi:10.1242/jeb.048967.

Link, A., Marimuthu, G., and Neuweiler, G. (1986). Movement as a specific stimulus for prey catching behaviour in rhinolophid and hipposiderid bats. J. Comp. Physiol. A 159, 403–413. doi:10.1007/BF00603985.

Mata, V. A., Rebelo, H., Amorim, F., McCracken, G. F., Jarman, S., and Beja, P. (2018). How much is enough? Effects of technical and biological replication on metabarcoding dietary analysis. Mol. Ecol. 0. doi:10.1111/mec.14779.

McCracken, G. F., Westbrook, J. K., Brown, V. A., Eldridge, M., Federico, P., and Kunz, T. H. (2012). Bats Track and Exploit Changes in Insect Pest Populations. PLoS ONE 7, e43839. doi:10.1371/journal.pone.0043839.

Milodowski, D. T., Coomes, D. A., Swinfield, T., Jucker, T., Riutta, T., Malhi, Y., et al. (2021). The impact of logging on vertical canopy structure across a gradient of tropical forest degradation intensity in Borneo. J. Appl. Ecol. doi:https://doi.org/10.1111/1365-2664.13895.

Neuweiler, G. (1990). Auditory adaptations for prey capture in echolocating bats. Physiol. Rev. 70, 615–641. doi:10.1152/physrev.1990.70.3.615.

Nielsen, A., and Bascompte, J. (2007). Ecological networks, nestedness and sampling effort. J. Ecol. 95, 1134–1141. doi:10.1111/j.1365-2745.2007.01271.x.

Novotný, V., and Basset, Y. (2000). Rare species in communities of tropical insect herbivores: pondering the mystery of singletons. Oikos 89, 564–572. doi:10.1034/j.1600-0706.2000.890316.x.

Oksanen, J., Blanchet, F. G., Friendly, M., Kindt, R., Legendre, P., McGlinn, D., et al. (2017). vegan: Community Ecology Package. Available at: https://CRAN.R-project.org/package=vegan.

de Oliveira, H. F. M., Camargo, N. F., Hemprich-Bennett, D. R., Rodríguez-Herrera, B., Rossiter, S. J., and Clare, E. L. (2020). Wing morphology predicts individual niche specialization in *Pteronotus mesoamericanus* (Mammalia: Chiroptera). PLOS ONE 15, e0232601. doi:10.1371/journal.pone.0232601.

R Core Team (2017). R: A Language and Environment for Statistical Computing. Vienna, Austria: R Foundation for Statistical Computing Available at: https://www.R-project.org/.

Ratnasingham, S., and Hebert, P. D. (2007). BOLD: The Barcode of Life Data System (http://www.barcodinglife.org). Mol. Ecol. Resour. 7, 355–364. doi:10.1111/j.1471-8286.2007.01678.x.

Reynolds, G., Payne, J., Sinun, W., Mosigil, G., and Walsh, R. P. D. (2011). Changes in forest land use and management in Sabah, Malaysian Borneo, 1990-2010, with a focus on the Danum Valley region. Philos. Trans. R. Soc. B Biol. Sci. 366, 3168–3176. doi:10.1098/rstb.2011.0154.

Rivera-Hutinel, A., Bustamante, R. O., Marín, V. H., and Medel, R. (2012). Effects of sampling completeness on the structure of plant–pollinator networks. Ecology 93, 1593–1603. doi:10.1890/11-1803.1.

Salinas-Ramos, V. B., Herrera Montalvo, L. G., León-Regagnon, V., Arrizabalaga-Escudero, A., and Clare, E. L. (2015). Dietary overlap and seasonality in three species of mormoopid bats from a tropical dry forest. Mol. Ecol. 24, 5296–5307. doi:10.1111/mec.13386.

Schloss, P. D., Westcott, S. L., Ryabin, T., Hall, J. R., Hartmann, M., Hollister, E. B., et al. (2009). Introducing mothur: Open-Source, Platform-Independent, Community-Supported Software for Describing and Comparing Microbial Communities. Appl. Environ. Microbiol. 75, 7537–7541. doi:10.1128/AEM.01541-09.

Schnitzler, H.-U., and Kalko, E. K. V. (2001). Echolocation by Insect-Eating Bats. BioScience 51, 557–569. doi:10.1641/0006-3568(2001)051[0557:EBIEB]2.0.CO;2.

Slade, E. M., Mann, D. J., and Lewis, O. T. (2011). Biodiversity and ecosystem function of tropical forest dung beetles under contrasting logging regimes. Biol. Conserv. 144, 166–174. doi:10.1016/j.biocon.2010.08.011.

Struebig, M. J., Kingston, T., Zubaid, A., Le Comber, S. C., Mohd-Adnan, A., Turner, A., et al. (2009). Conservation importance of limestone karst outcrops for Palaeotropical bats in a fragmented landscape. Biol. Conserv. 142, 2089–2096. doi:10.1016/j.biocon.2009.04.005.

Struebig, M. J., Turner, A., Giles, E., Lasmana, F., Tollington, S., Bernard, H., et al. (2013). Quantifying the Biodiversity Value of Repeatedly Logged Rainforests. Adv. Ecol. Res. 48, 183–224. doi:10.1016/B978-0-12-417199-2.00003-3.

Tournayre, O., Leuchtmann, M., Galan, M., Trillat, M., Piry, S., Pinaud, D., et al. (2021). eDNA metabarcoding reveals a core and secondary diets of the greater horseshoe bat with strong spatio-temporal plasticity. Environ. DNA 3, 277–296. doi:https://doi.org/10.1002/edn3.167.

Vallejo, N., Aihartza, J., Goiti, U., Arrizabalaga-Escudero, A., Flaquer, C., Puig, X., et al. (2019). The diet of the notch-eared bat (*Myotis emarginatus*) across the Iberian Peninsula analysed by amplicon metabarcoding. Hystrix Ital. J. Mammal. 30, 59–64. doi:10.4404/hystrix-00189-2019.

Wei, T., and Simko, V. (2017). R package “corrplot”: Visualization of a correlation matrix. Available at: https://github.com/taiyun/corrplot.

Zeale, M. R. K., Butlin, R. K., Barker, G. L. A., Lees, D. C., and Jones, G. (2011). Taxon-specific PCR for DNA barcoding arthropod prey in bat faeces. Mol. Ecol. Resour. 11, 236–244. doi:10.1111/j.1755-0998.2010.02920.x.

