## Supplementary information for "Selective logging shows no impact on the dietary breadth of the fawn leaf-nosed bat (*Hipposideros cervinus*)"

### Supplementary table 1: Map of study sites

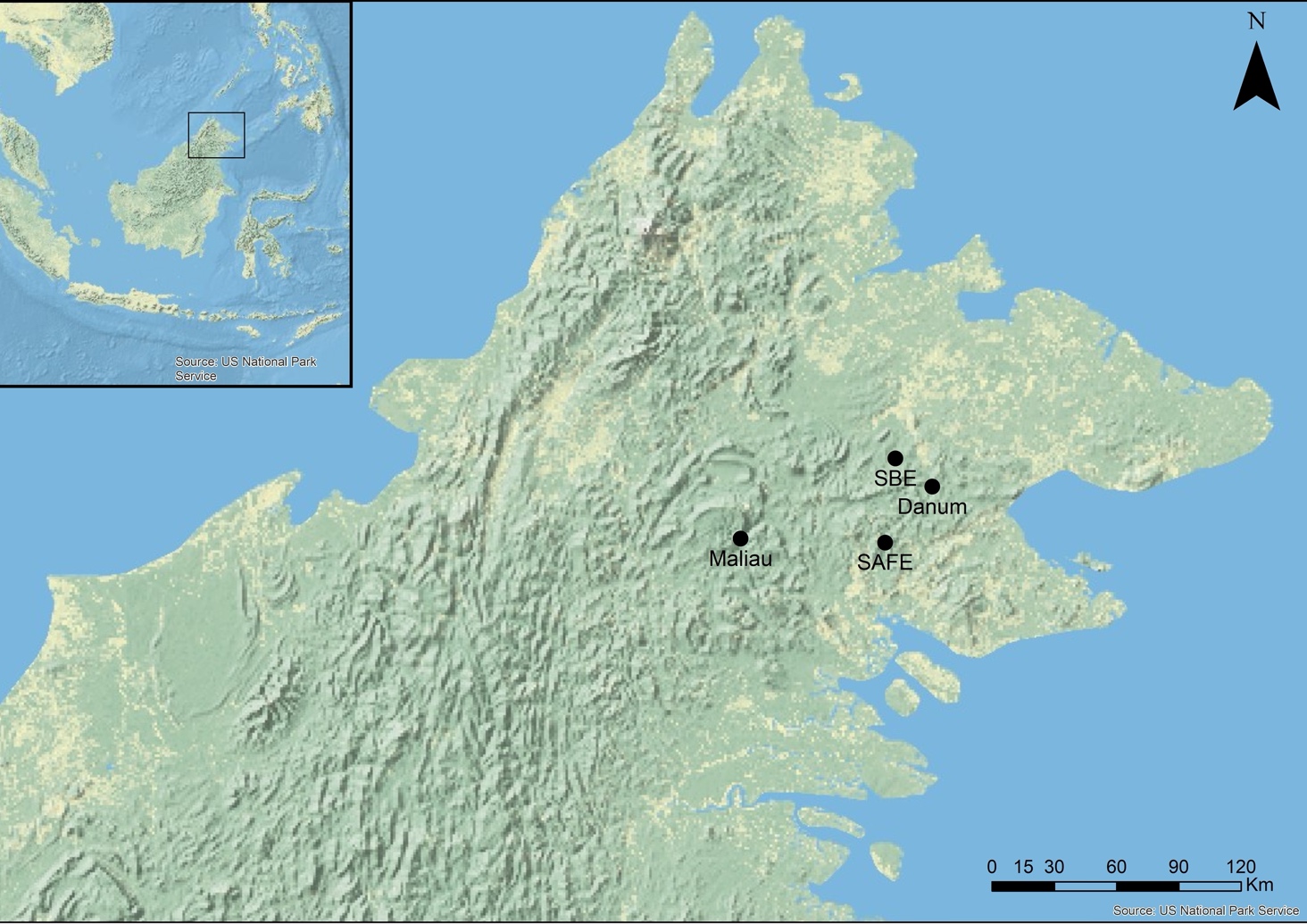

### Supplementary information 1: MOTU richness at different clustering levels and QC parameters

| **Site** | **clustering** | **Post-clustering QC** | **Observed richness** | **Estimated richness** | **% completeness** | **No. samples** | **Estimated no. samples required** | **Standard error** | **Lower 95% confidence level** | **Upper 95% confidence level** |
| --- | --- | --- | --- | --- | --- | --- | --- | --- | --- | --- |
| SAFE | 91 | None | 1113 | 3742.4 | 29.7 | 111 | 373.2 | 289.2 | 3233.9 | 4372.8 |
| Maliau | 91 | None | 975 | 2118.1 | 46.0 | 97 | 210.7 | 130.0 | 1890.4 | 2402.5 |
| SBE | 91 | None | 595 | 1667.9 | 35.7 | 57 | 159.8 | 162.7 | 1393.3 | 2036.8 |
| Danum | 91 | None | 2119 | 4749.3 | 44.6 | 186 | 416.9 | 208.8 | 4370.8 | 5191.5 |
| SAFE | 92 | None | 1291 | 4817.1 | 26.8 | 111 | 414.2 | 360.2 | 4178.7 | 5596.5 |
| Maliau | 92 | None | 1139 | 2974.6 | 38.3 | 97 | 253.3 | 194.1 | 2631.9 | 3395.9 |
| SBE | 92 | None | 661 | 1923.8 | 34.4 | 57 | 165.9 | 179.6 | 1618.0 | 2327.3 |
| Danum | 92 | None | 2611 | 6879.6 | 38.0 | 186 | 490.1 | 304.3 | 6323.7 | 7518.7 |
| SAFE | 93 | None | 1526 | 7727.8 | 19.7 | 111 | 562.1 | 632.1 | 6607.4 | 9095.1 |
| Maliau | 93 | None | 1389 | 4716.2 | 29.5 | 97 | 329.4 | 325.6 | 4136.8 | 5417.8 |
| SBE | 93 | None | 761 | 2605.9 | 29.2 | 57 | 195.2 | 248.0 | 2180.3 | 3159.1 |
| Danum | 93 | None | 3336 | 10300.6 | 32.4 | 186 | 574.3 | 439.3 | 9491.4 | 11216.2 |
| SAFE | 94 | None | 1680 | 9187.9 | 18.3 | 111 | 607.1 | 744.9 | 7864.1 | 10795.2 |
| Maliau | 94 | None | 1559 | 5711.3 | 27.3 | 97 | 355.4 | 388.3 | 5017.3 | 6544.6 |
| SBE | 94 | None | 917 | 3244.9 | 28.3 | 57 | 201.7 | 285.7 | 2748.9 | 3875.3 |
| Danum | 94 | None | 3875 | 13582.5 | 28.5 | 186 | 652.0 | 580.4 | 12509.9 | 14788.3 |

| **Site** | **clustering** | **Post-clustering QC** | **Observed richness** | **Estimated richness** | **% completeness** | **No. samples** | **Estimated no. samples required** | **Standard error** | **Lower 95% confidence level** | **Upper 95% confidence level** |
| --- | --- | --- | --- | --- | --- | --- | --- | --- | --- | --- |
| SAFE | 95 | None | 2072 | 14895.9 | 13.9 | 111 | 798.0 | 1250.0 | 12670.5 | 17588.6 |
| Maliau | 95 | None | 1958 | 8513.8 | 23.0 | 97 | 421.8 | 565.0 | 7496.5 | 9717.8 |
| SBE | 95 | None | 1162 | 5393.6 | 21.5 | 57 | 264.6 | 492.4 | 4533.3 | 6473.5 |
| Danum | 95 | None | 5142 | 23249.2 | 22.1 | 186 | 841.0 | 989.7 | 21410.9 | 25295.1 |
| SAFE | 96 | None | 2397 | 19854.3 | 12.1 | 111 | 919.4 | 1666.9 | 16880.8 | 23438.3 |
| Maliau | 96 | None | 2273 | 11216.1 | 20.3 | 97 | 478.6 | 739.9 | 9879.5 | 12787.6 |
| SBE | 96 | None | 1389 | 6707.6 | 20.7 | 57 | 275.3 | 571.6 | 5700.1 | 7950.6 |
| Danum | 96 | None | 6168 | 30818.6 | 20.0 | 18 | 929.4 | 1258.1 | 28473.4 | 33410.3 |
| SAFE | 97 | None | 3555 | 42042.1 | 8.5 | 111 | 1312.7 | 3466.9 | 35824.6 | 49457.6 |
| Maliau | 97 | None | 3335 | 21441.8 | 15.6 | 97 | 623.6 | 1341.9 | 18996.9 | 24268.3 |
| SBE | 97 | None | 2047 | 11649.1 | 17.6 | 57 | 324.4 | 888.9 | 10058.8 | 13554.9 |
| Danum | 97 | None | 9479 | 58971.7 | 16.1 | 186 | 1157.2 | 2161.2 | 54913.9 | 63392.0 |
| SAFE | 98 | None | 4667 | 64442.8 | 7.2 | 111 | 1532.7 | 5006.0 | 55408.6 | 75085.5 |
| Maliau | 98 | None | 4448 | 36192.4 | 12.3 | 97 | 789.3 | 2212.4 | 32144.0 | 40832.7 |
| SBE | 98 | None | 2839 | 20418.0 | 13.9 | 57 | 409.9 | 1490.4 | 17731.0 | 23589.7 |
| Danum | 98 | None | 13115 | 102587.5 | 12.8 | 186 | 1454.9 | 3584.8 | 95832.8 | 109893.8 |
| SAFE | 91 | LULU | 823 | 2179.7 | 37.8 | 111 | 294.0 | 167.8 | 1888.5 | 2550.4 |
| Maliau | 91 | LULU | 770 | 1422.3 | 54.1 | 97 | 179.2 | 84.6 | 1276.3 | 1610.2 |
| SBE | 91 | LULU | 467 | 1162.5 | 40.2 | 57 | 141.9 | 119.2 | 965.4 | 1437.7 |
| Danum | 91 | LULU | 1422 | 2559.1 | 55.6 | 186 | 334.7 | 112.9 | 2358.4 | 2802.8 |

| **Site** | **clustering** | **Post-clustering QC** | **Observed richness** | **Estimated richness** | **% completeness** | **No. samples** | | **Estimated no. samples required** | **Standard error** | **Lower 95% confidence level** | | **Upper 95% confidence level** |
| --- | --- | --- | --- | --- | --- | --- | --- | --- | --- | --- | --- | --- |
| SAFE | 92 | LULU | 819 | 2252.6 | 36.4 | 111 | | 305.3 | 174.1 | 1950.0 | | 2636.3 |
| Maliau | 92 | LULU | 751 | 1488.3 | 50.5 | 97 | | 192.2 | 95.3 | 1323.9 | | 1699.9 |
| SBE | 92 | LULU | 450 | 1047.6 | 43.0 | 57 | 132.7 | | 101.4 | | 879.5 | 1281.4 |
| Danum | 92 | LULU | 1451 | 2750.6 | 52.8 | 186 | 352.6 | | 124.9 | | 2528.0 | 3019.3 |
| SAFE | 93 | LULU | 784 | 2546.9 | 30.8 | 111 | 360.6 | | 225.7 | | 2157.0 | 3047.5 |
| Maliau | 93 | LULU | 769 | 1696.9 | 45.3 | 97 | 214.0 | | 116.9 | | 1494.5 | 1955.6 |
| SBE | 93 | LULU | 437 | 1155.7 | 37.8 | 57 | 150.7 | | 124.7 | | 949.8 | 1444.2 |
| Danum | 93 | LULU | 1454 | 2834.9 | 51.3 | 186 | 362.6 | | 130.2 | | 2602.3 | 3114.5 |
| SAFE | 94 | LULU | 758 | 2397.4 | 31.6 | 111 | 351.1 | | 211.7 | | 2032.1 | 2867.4 |
| Maliau | 94 | LULU | 770 | 1755.0 | 43.9 | 97 | 221.1 | | 123.9 | | 1540.6 | 2029.1 |
| SBE | 94 | LULU | 447 | 1086.4 | 41.1 | 57 | 138.5 | | 108.5 | | 906.5 | 1336.6 |
| Danum | 94 | LULU | 1455 | 2975.5 | 48.9 | 186 | 380.4 | | 143.8 | | 2718.7 | 3284.3 |
| SAFE | 95 | LULU | 758 | 2894.8 | 26.2 | 111 | 423.9 | | 286.7 | | 2402.7 | 3534.3 |
| Maliau | 95 | LULU | 744 | 1723.4 | 43.2 | 97 | 224.7 | | 124.9 | | 1507.7 | 2000.2 |
| SBE | 95 | LULU | 442 | 1186.5 | 37.3 | 57 | 153.0 | | 129.3 | | 973.0 | 1485.7 |
| Danum | 95 | LULU | 1476 | 3183.5 | 46.4 | 186 | 401.2 | | 159.2 | | 2898.9 | 3524.9 |
| SAFE | 96 | LULU | 754 | 2849.9 | 26.5 | 111 | 419.5 | | 281.7 | | 2366.3 | 3478.5 |
| Maliau | 96 | LULU | 727 | 1671.7 | 43.5 | 97 | 223.0 | | 121.7 | | 1461.6 | 1941.9 |
| SBE | 96 | LULU | 440 | 1059.0 | 41.5 | 57 | 137.2 | | 106.1 | | 883.5 | 1304.0 |
| Danum | 96 | LULU | 1487 | 3314.4 | 44.9 | 186 | 414.6 | | 169.5 | | 3011.3 | 3677.8 |

| **Site** | **clustering** | **Post-clustering QC** | **Observed richness** | **Estimated richness** | **% completeness** | **No. samples** | **Estimated no. samples required** | **Standard error** | **Lower 95 % confidence level** | **Upper 95% confidence level** |
| --- | --- | --- | --- | --- | --- | --- | --- | --- | --- | --- |
| SAFE | 97 | LULU | 780 | 3020.6 | 25.8 | 111 | 429.9 | 296.6 | 2510.6 | 3681.0 |
| Maliau | 97 | LULU | 741 | 1762.4 | 42.0 | 97 | 230.7 | 130.8 | 1536.5 | 2052.4 |
| SBE | 97 | LULU | 451 | 1124.5 | 40.1 | 57 | 142.1 | 113.5 | 936.2 | 1385.9 |
| Danum | 97 | LULU | 1555 | 3463.9 | 44.9 | 186 | 414.3 | 170.7 | 3157.6 | 3828.7 |
| SAFE | 98 | LULU | 807 | 3328.6 | 24.2 | 111 | 457.8 | 334.3 | 2753.8 | 4073.2 |
| Maliau | 98 | LULU | 760 | 1891.5 | 40.2 | 97 | 241.4 | 143.5 | 1643.3 | 2209.4 |
| SBE | 98 | LULU | 490 | 1260.6 | 38.9 | 57 | 146.6 | 124.9 | 1051.9 | 1546.7 |
| Danum | 98 | LULU | 1649 | 3644.2 | 45.2 | 186 | 411.1 | 171.4 | 3335.6 | 4009.3 |

s

### Supplementary information 2: Family-level diet of *H. cervinus*

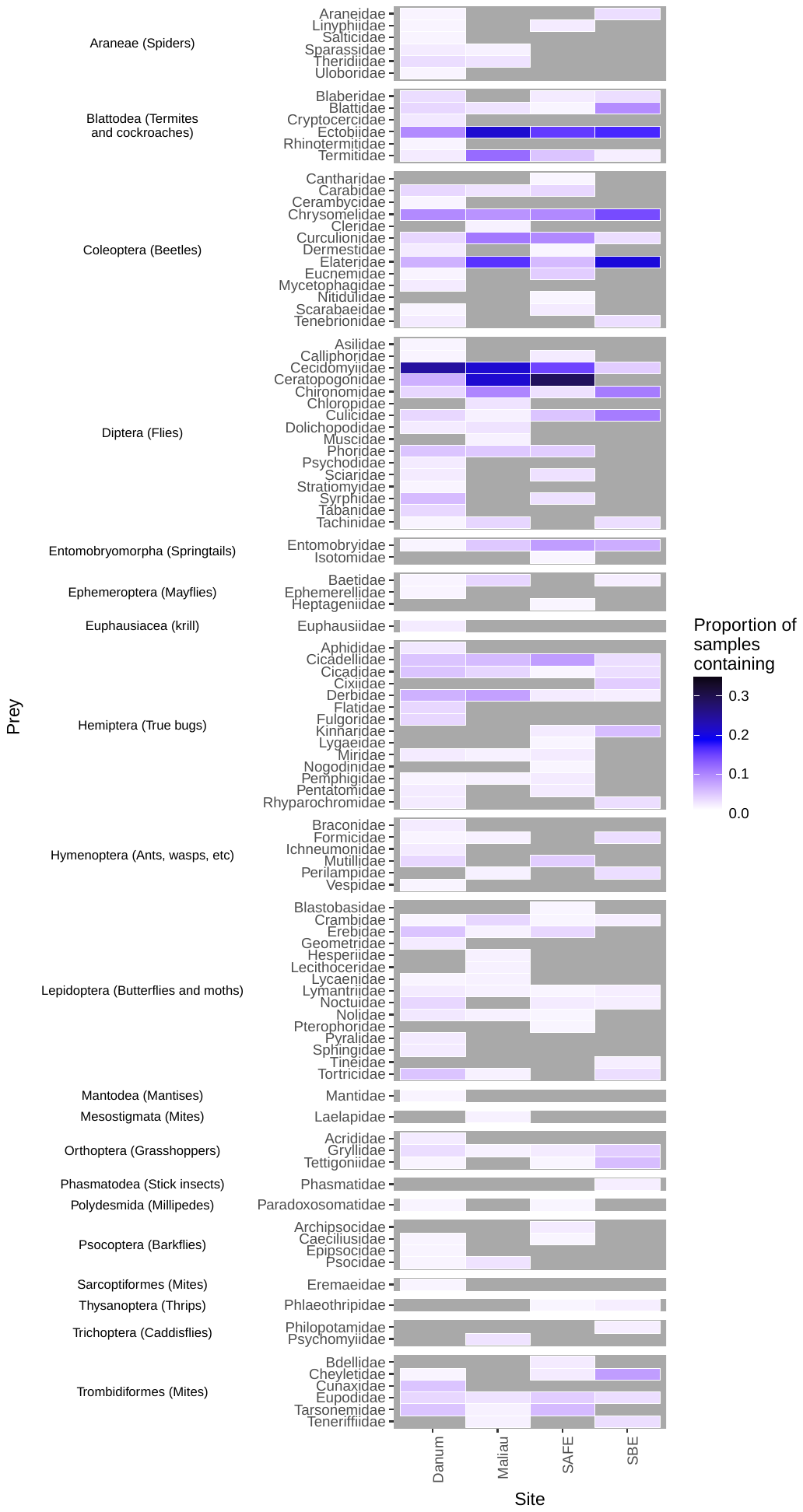

### Supplementary information 3: Correlations between prey orders consumed by *H. cervinus*

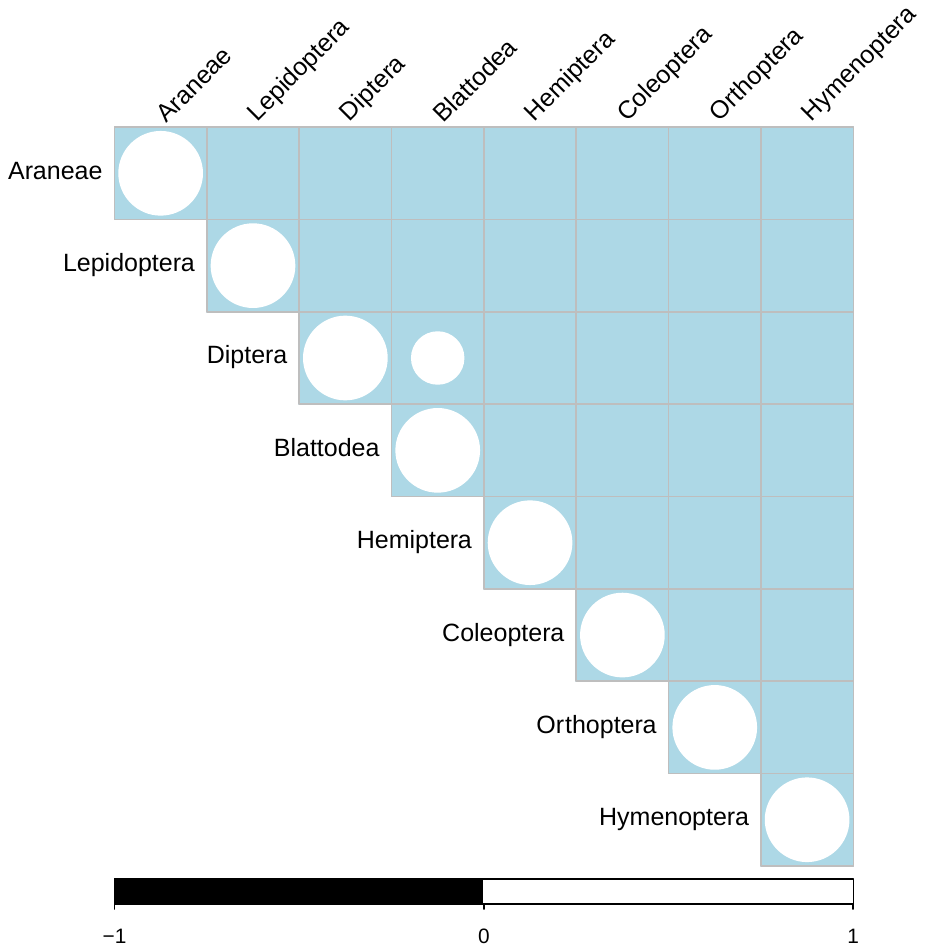

**Figure 4.3. Positive and negative correlations of orders consumed by bats.** Circle size is proportional to the strength of the correlation, black circles are negative correlations, white circles are positive correlations. Empty locations within the grid show non-significant (p>0.05) correlations. The plot shows, for example, a negative correlation between the presence of Araneae in a bat’s diet and the presence of Hymenoptera, but a positive correlation between the presence of Diptera and Hymenoptera.

### Supplementary information 4: Significant values from SIMPER analysis, rounded to three decimal places

| Order | Sites | Taxa | average | sd | ratio | ava | avb | cumsum | p |
| --- | --- | --- | --- | --- | --- | --- | --- | --- | --- |
| Diptera | SAFE-Maliau | Diptera | 0.151 | 0.147 | 1.027 | 1.29 | 1.484 | 0.5 | 0.003 |
| Coleoptera | SAFE-Maliau | Coleoptera | 0.091 | 0.107 | 0.855 | 0.524 | 0.912 | 0.792 | 0.003 |
| Psocoptera | SAFE-Maliau | Psocoptera | 0.011 | 0.041 | 0.264 | 0.065 | 0.033 | 0.963 | 0.01 |
| Mesostigmata | SAFE-Maliau | Mesostigmata | 0.001 | 0.008 | 0.102 | 0 | 0.011 | 0.998 | 0.005 |
| Blattodea | SAFE-SBE | Blattodea | 0.172 | 0.18 | 0.955 | 0.734 | 2.56 | 0.507 | 0.014 |
| Orthoptera | SAFE-SBE | Orthoptera | 0.032 | 0.073 | 0.441 | 0.185 | 0.3 | 0.882 | 0.03 |
| Thysanoptera | SAFE-SBE | Thysanoptera | 0.009 | 0.04 | 0.223 | 0.024 | 0.04 | 0.97 | 0.02 |
| Cyclopoida | SAFE-SBE | Cyclopoida | 0.005 | 0.03 | 0.173 | 0.008 | 0.04 | 0.986 | 0.034 |
| Blattodea1 | Maliau-SBE | Blattodea | 0.175 | 0.185 | 0.946 | 1.242 | 2.56 | 0.52 | 0.014 |
| Trichoptera | Maliau-SBE | Trichoptera | 0.003 | 0.015 | 0.195 | 0.022 | 0.02 | 0.997 | 0.029 |
| Ephemeroptera | Maliau-Danum | Ephemeroptera | 0.004 | 0.025 | 0.177 | 0.033 | 0.011 | 0.987 | 0.035 |
| Blattodea2 | SBE-Danum | Blattodea | 0.18 | 0.191 | 0.945 | 2.56 | 1.603 | 0.547 | 0.003 |
| Cyclopoida1 | SBE-Danum | Cyclopoida | 0.005 | 0.029 | 0.176 | 0.04 | 0.011 | 0.979 | 0.028 |
| Phasmatodea | SBE-Danum | Phasmatodea | 0.004 | 0.029 | 0.138 | 0.02 | 0.006 | 0.99 | 0.046 |
